# The processing of dynamic faces in the human brain: Support for an integrated neural framework of face processing

**DOI:** 10.1101/140939

**Authors:** Michal Bernstein, Yaara Erez, Idan Blank, Galit Yovel

**Affiliations:** Sagol School of Neuroscience, Tel Aviv University, Tel Aviv 6997801, Israel; MRC Cognition and Brain Sciences Unit, 15 Chaucer Rd, Cambridge, UK; Department of Brain and Cognitive Sciences, Massachusetts Institute of Technology, Cambridge, MA; School of Psychological Sciences, Tel Aviv University, Tel Aviv 6997801, Israel

**Keywords:** Face perception, Motion, FFA, Superior temporal sulcus (STS), MT

## Abstract

Faces convey rich information including identity, gender and expression. Current neural models of face processing suggest a dissociation between the processing of invariant facial aspects such as identity and gender, that engage the fusiform face area (FFA) and the processing of changeable aspects, such as expression and eye gaze, that engage the posterior superior temporal sulcus face area (pSTS-FA). Recent studies report a second dissociation within this network such that the pSTS-FA, but not the FFA, shows much stronger response to dynamic than static faces. The aim of the current study was to test a unified model that accounts for these two major functional characteristics of the neural face network. In an fMRI experiment, we presented static and dynamic faces while subjects judged an invariant (gender) or a changeable facial aspect (expression). We found that the pSTS-FA was more engaged in processing dynamic than static faces and changeable than invariant facial aspects, whereas the OFA and FFA showed similar response across all four conditions. Our results reveal no dissociation between the processing of changeable and invariant facial aspects, but higher sensitivity to the processing of changeable facial aspects by the motion-sensitive face area in the superior temporal sulcus.

---

Faces engage a set of localized cortical regions in the occipital temporal cortex, forming a neural network that is specialized for face processing (for reviews see Kanwisher and Yovel, 2006, Duchaine and Yovel, 2015). According to a dominant neural model of face processing (Haxby et al., 2000, Gobbini and Haxby, 2007), this network is composed of three core areas: the occipital face area (OFA), the fusiform face area (FFA) and the posterior superior temporal sulcus face area (pSTS-FA). The model further posits that these core areas are divided into two pathways that play different functional roles: a ventral pathway, including the FFA, processes invariant facial aspects, such as identity and gender, whereas a dorsal pathway, including the pSTS-FA, processes changeable facial aspects such as facial expression and eye gaze. The OFA, according to the model, is involved in early stages of processing and provides input to both pathways (see Figure 1A).

**Figure 1.**
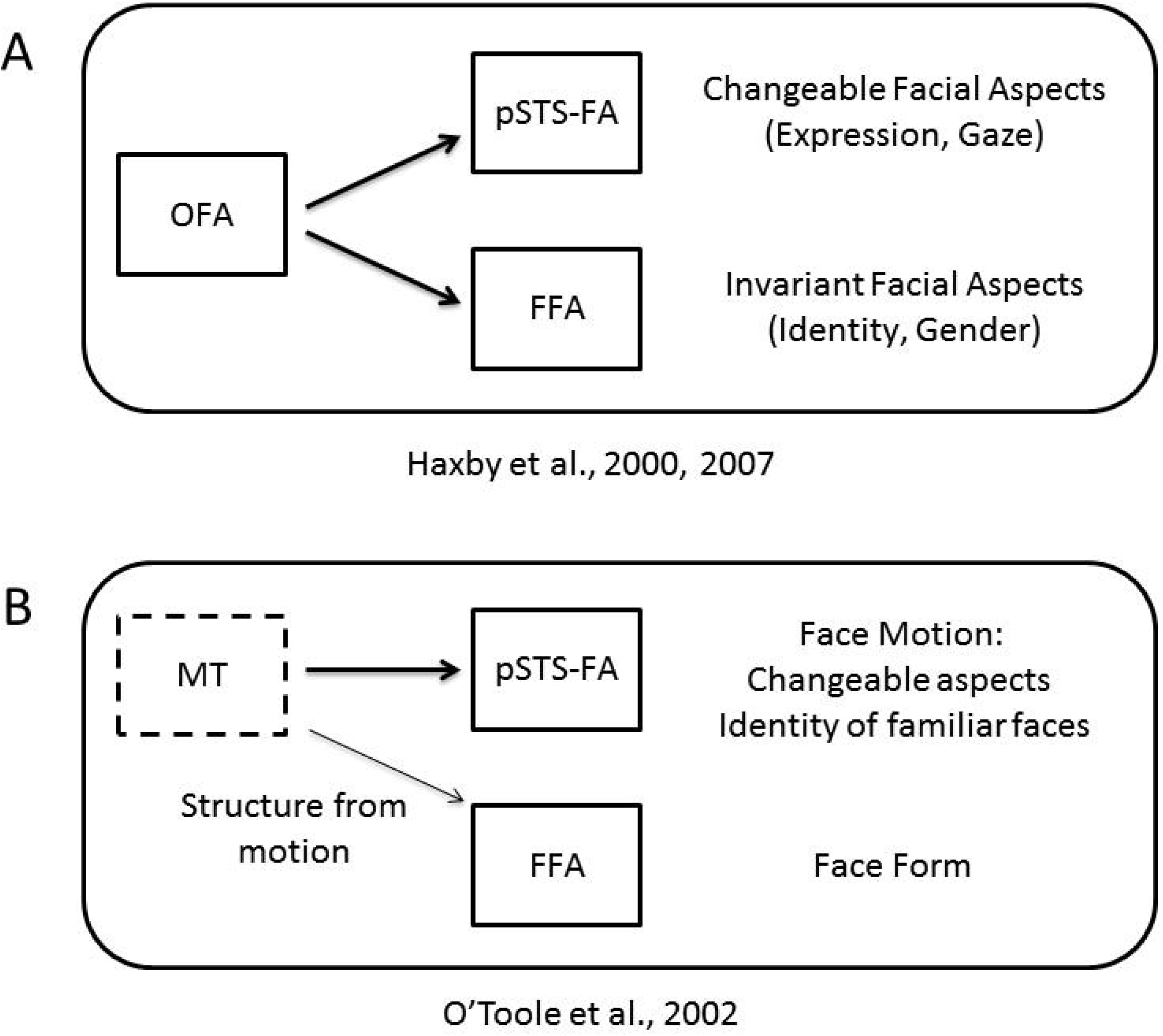
A. The current dominant neural model of face processing (Haxby et al., 2000, Gobbini and Haxby, 2007) includes two pathways: a ventral pathway, including the FFA, processes invariant facial aspects, such as identity and gender, whereas a dorsal pathway, including the pSTS-FA, processes changeable facial aspects such as facial expression and eye gaze. The OFA is involved in early stages of face processing and provides input to both pathways. B. Modifications to this model proposed by (O’Toole et al., 2002) for dynamic faces: the ventral pathway processes facial form while the dorsal pathway processes facial motion. The motion-selective area MT provides input to the dorsal, motion pathway, and may also send “motionless” structure information to the ventral, form pathway.

Many studies have provided evidence to support the role of the FFA in the processing of face identity (Grill-Spector et al., 2004, Rotshtein et al., 2005, Yovel and Kanwisher, 2005, Gilaie-Dotan and Malach, 2007, Nestor et al., 2011, Baseler et al., 2013, Goesaert and Op de Beeck, 2013, Axelrod and Yovel, 2015) and the role of the pSTS-FA in the processing of changeable facial aspects, and facial expression in particular (Engell and Haxby, 2007, Furl et al., 2007, Schultz and Pilz, 2009, Said et al., 2010). However, numerous studies have also reported similar sensitivity to facial expression in the FFA as in the pSTS-FA (Ganel et al., 2005, Fox et al., 2009b, Kadosh et al., 2010, Xu and Biederman, 2010; for a recent review see Bernstein and Yovel, 2015), challenging the prevalent view that the FFA processes invariant but not changeable facial aspects (Haxby et al., 2000, Gobbini and Haxby, 2007). Importantly, all these studies used static face images, whereas in real life faces are typically seen in motion.

Recent neuroimaging studies that have examined the processing of dynamic faces (i.e. video-clips of moving face) revealed that the pSTS-FA shows a much stronger response to dynamic than static faces, whereas the FFA and OFA show similar responses to both dynamic and static faces (Fox et al., 2009a, Pitcher et al., 2011). A face-selective area in the inferior frontal gyrus (IFG-FA) was also reported in these studies to show stronger response to dynamic faces. These findings thus suggest that the dorsal face areas may be sensitive to facial motion while the ventral areas may not be sensitive to dynamic information from faces, but rather process facial form. These findings are consistent with a two stream model suggested by O’Toole and colleagues (O’Toole et al., 2002) in which the pSTS-FA extracts dynamic information from faces and the FFA extracts form information from both dynamic and static faces (Figure 1B). The idea of dissociable pathways for dynamic and static face information is further supported by a recent study (Pitcher et al., 2014), in which fMRI was combined with transcranial magnetic stimulation (TMS). Stimulation of the pSTS-FA resulted in a reduced fMRI signal to dynamic but not static faces, whereas stimulation of the OFA reduced the fMRI signal to static but not dynamic faces (Pitcher et al., 2014). Finally, fMRI studies on monkeys also show that the dorsal but not the ventral face patches in the macaque’s STS are sensitive to facial motion (Fisher and Freiwald, 2015, for reviews see Yovel and Freiwald, 2013, Freiwald et al., 2016).

Taking into consideration the dynamic nature of changeable facial aspects, these recent findings of sensitivity in the pSTS-FA to dynamic information may fit with the notion that this area is engaged in processing changeable facial aspects. Changeable aspects, such as facial expression, depend on the dynamic changes of the face structure over time. Indeed, behavioral studies have consistently shown a ‘dynamic advantage’ for expression recognition, meaning that expressions are better recognized when presented in motion (Bassili, 1978, 1979, Ambadar et al., 2005, Nusseck et al., 2008, Trautmann et al., 2009, Fujimura and Suzuki, 2010, Kaulard et al., 2012, Ceccarini and Caudek, 2013; for a review see Krumhuber et al., 2013). In contrast, a dynamic advantage for invariant facial aspects such as identity is less conclusive and depends on stimulus and task factors: Studies have found that motion may serve as a useful supplementary cue for face recognition especially for familiar faces (O’Toole, 2010). For unfamiliar faces, the contribution of motion to face recognition may be limited to tasks that include dynamic information both at learning and at test (Roark et al., 2006, Lander and Davies, 2007) and when the viewing conditions are suboptimal (e.g. poor illumination, degraded stimuli) (for reviews see O’Toole, 2010, Xiao et al., 2014, Lander and Butcher, 2015).

Taken together, dorsal and ventral regions in the face-processing network show distinct patterns of sensitivity to changeable vs. invariant facial aspects as well as dynamic (motion) vs. static (form) information. In order to integrate these two related functional characterizations of the network, we have recently proposed a comprehensive neural model of face processing (Figure 2) (Bernstein and Yovel, 2015, Duchaine and Yovel, 2015). According to this model, the dorsal face areas are engaged in the processing of facial motion, and also in the processing of changeable facial aspects because they rely on dynamic information. In contrast, the ventral face areas are engaged in the processing of facial form, and thus are not sensitive to motion information and are recruited when processing both changeable and invariant facial aspects. Furthermore, based on previous findings of structural and functional connectivity among the face areas (Gschwind et al., 2012, Pyles et al., 2013, Avidan et al., 2014) we suggested that the OFA is primarily connected to the FFA but not the pSTS-FA. Finally, the IFG-FA, which is also sensitive to dynamic faces, was suggested to be part of the dorsal face pathway.

**Figure 2.**
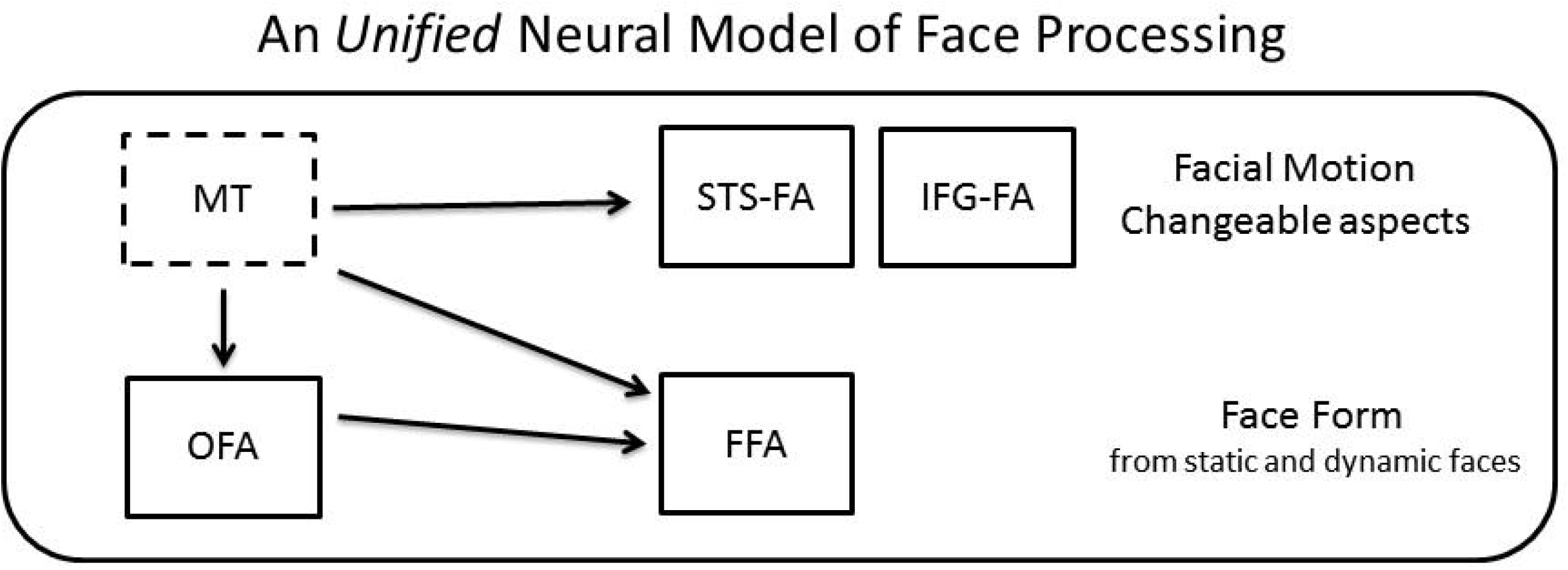
The unified neural model of face processing. The dorsal face areas are engaged in the processing of facial motion and changeable facial aspects, whereas the ventral areas are engaged in the processing of facial form. Area MT sends input to the dorsal areas for motion processing, and to the ventral areas for structure-from-motion analysis.

Here, we provide the first direct test of this model. To this end, we used a factorial design that crossed task requirements for processing different facial aspects with stimuli that either contained motion information in them or did not. Specifically, subjects performed alternate categorization tasks that focused on an invariant facial aspect (i.e., gender) or a changeable facial aspect (i.e., expression) on both static and dynamic faces. We assessed the fMRI responses to these different tasks and stimuli classes in the ventral and dorsal face areas, as well as the motion area, MT. This design allowed us to test the following three hypotheses regarding the division of labor between the dorsal and ventral face areas (see Figure 3): (i) if the primary division of labor is to changeable vs. invariant facial aspects (Figure 3A), then the pSTS-FA should respond strongly during the expression task and the FFA during the gender task, for both static and dynamic faces; (ii) if the primary division of labor is to motion vs. form processing (Figure 3B), then the pSTS-FA should be responsive to dynamic faces more than static whereas the FFA should show similar response to both dynamic and static faces, regardless of the task; (iii) Based on our integrated model, the pSTS-FA would show a larger response to dynamic than static faces, and also during the expression than the gender task, whereas the FFA would be similarly sensitive to dynamic and static faces and to changeable and invariant facial aspects (Figure 3C). Finally, to assess the proposed links between the face areas for the processing of dynamic faces, we also examined the functional connectivity among the face areas and with the motion area, MT.

**Figure 3.**
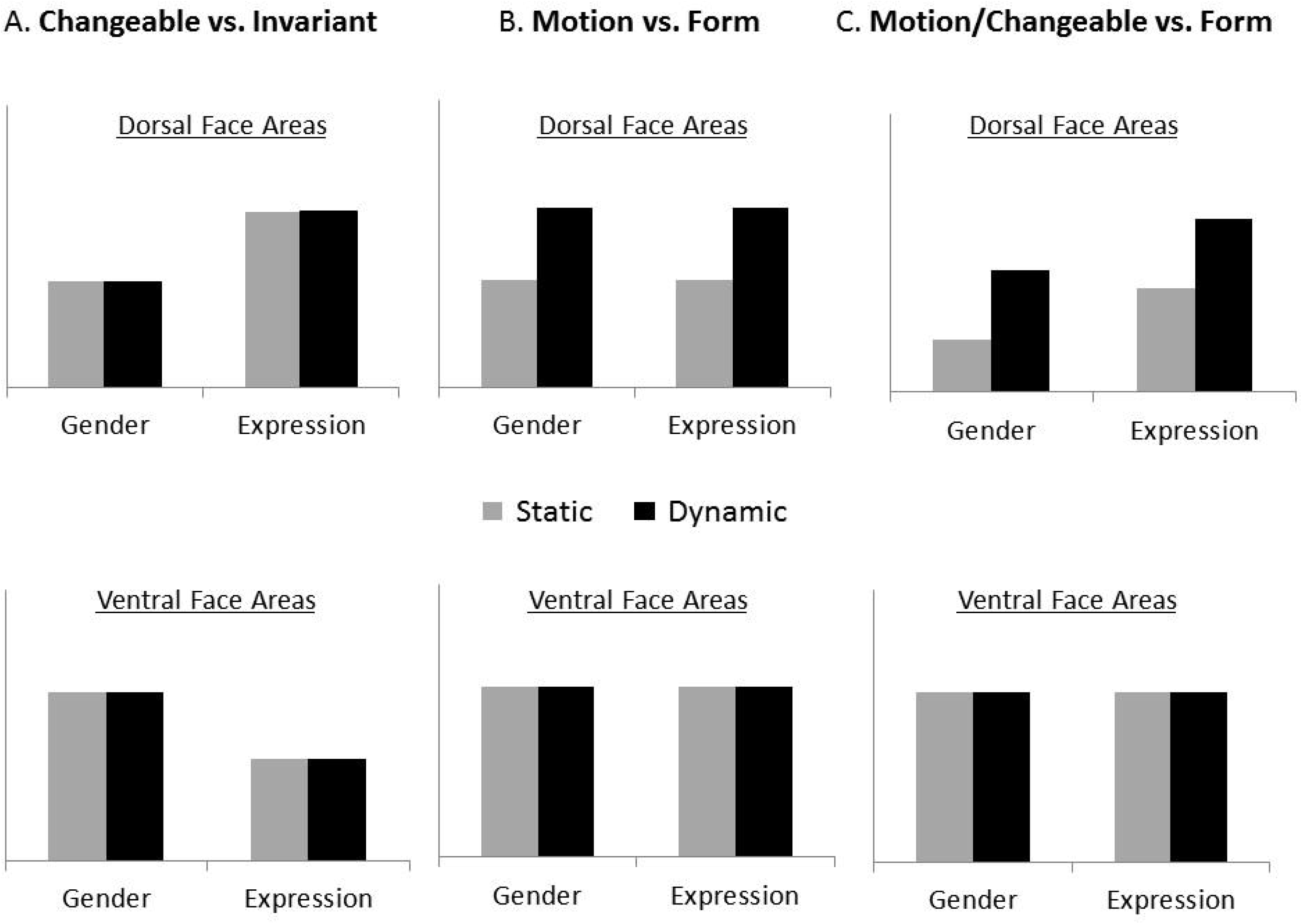
Three alternative predictions were tested: A. The primary division of labor between the dorsal and ventral face areas is to changeable and invariant aspects, as proposed by the current face processing neural model (Haxby et al., 2000, Gobbini and Haxby, 2007). B. The primary division of labor is to motion and form processing. C. An integrated model in which the dorsal face areas are sensitive to both face motion and changeable facial aspects and the ventral face areas extract form information from dynamic and static faces for both invariant and changeable aspects.

## Experiment 1

In Experiment 1 we compared the fMRI response of face-selective areas to dynamic and static faces while participants performed an expression (i.e. changeable) or gender (i.e. invariant) categorization task. In particular, participants had to judge whether a face has a positive or negative facial expression or whether it was male or female. The static face stimuli were four static images taken from the dynamic face videos (see Figure 4). Such multi-static images provide richer information than one static image and therefore better match the dynamic stimuli (e.g., O’Toole et al., 2011). The multi-static stimuli were presented in the order of their presentation in the video. To minimize apparent motion, we selected one image per second.

**Figure 4.**
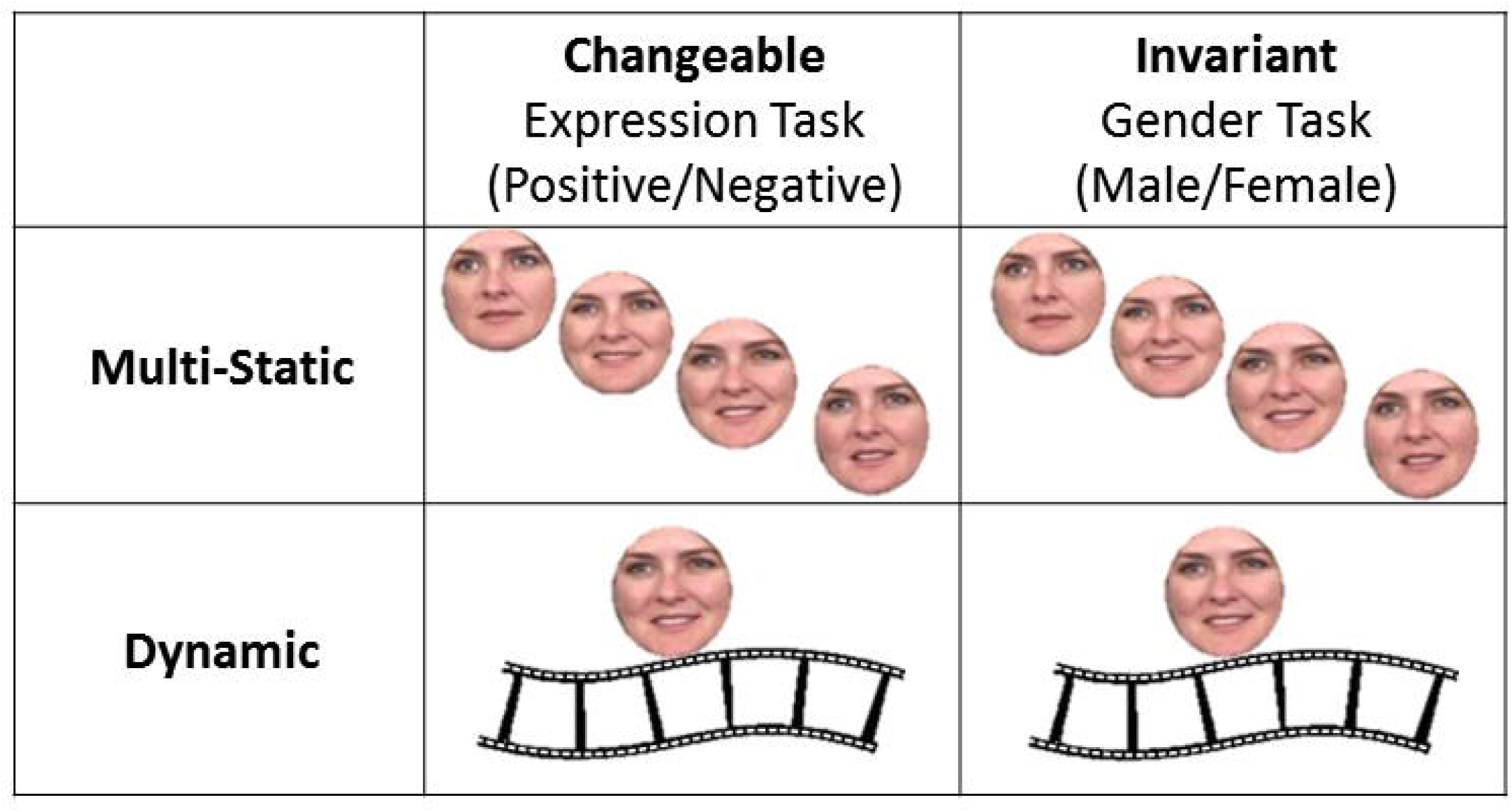
Faces from the database of moving and static faces collected at the Vision Lab at UTD (O’Toole et al., 2005) were cropped to show only the internal facial features, with no hair or other external facial features. The dynamic conditions included 4 second movie-clips of neutral faces turning happy or disgust. The static conditions included four images from the movie (the first frame of each second) presented for 1 second each in order of appearance in the movie (Experiment 1) or in a scrambled order (Experiment 2).

### Materials and Methods

#### Participants

Twenty healthy volunteers (8 men, ages 19–35) with normal or corrected-to-normal vision participated in the experiment. Two participants were excluded due to excessive head movements leaving a total of eighteen participants in the final analysis. Participants were paid $15/hr. All participants provided written informed consent to participate in the study, which was approved by the ethics committees of the Sheba Medical Center and Tel Aviv University.

#### Stimuli

The experiment included three parts: a functional dynamic face localizer to localize the face-selective areas, a functional motion localizer to localize the motion-selective area MT, and the main experiment.

##### Functional face localizer

Stimuli of the face localizer were dynamic (1s movie-clips) faces and objects (e.g., a ball, a fan). The object stimuli were provided by (Fox et al., 2009a). The face stimuli were filmed in our lab and included 16 different faces which moved their face/head by talking or making various facial expressions or mouth movements.

##### Functional motion localizer

Stimuli of the MT localizer were white dots moving on a black background, and static shots of these dots (Huk et al., 2002).

##### Main Experiment

Stimuli for the main experiment were taken from the database of moving and static faces collected at the Vision Lab at University Texas at Dallas (O’Toole et al., 2005). All stimuli were cropped to show only the face area, with no hair or other external facial features (see Figure 4 for example stimuli). The main experiment included four conditions in a factorial design that crossed stimulus class (dynamic or static) and categorization task (expression or gender): Dynamic faces-Expression task, Static faces-Expression task, Dynamic faces-Gender task, and Static faces-Gender task. In the dynamic conditions, 4s movie-clips (24 frames per second) were presented. In the static conditions, four images from the movie (the first frame of each second) were presented for 1 second each, in their chronological order of appearance within the movie. 48 different identities were used as stimuli, half females and half males. Twenty-seven of the movies (12 females, 15 males) showed a face transitioning from a neutral expression to a happy (positive) expression, and the remaining 21 movies showed a face transitioning from a neutral expression to a disgust (negative) expression. Importantly, the same 48 identities were used in all four experimental conditions, but the stimuli were counterbalanced across subjects such that each subject saw a given identity in only one of the four conditions.

#### Apparatus and Procedure

fMRI data were acquired in a 3T Siemens MAGNETOM Prisma MRI scanner, using a 64-channel head coil. Echo-planar volumes were acquired with the following parameters: repetition time (TR) = 2s, echo time = 35ms, flip angel = 90°, 31 slices per TR, slice thickness = 3mm, field of view = 20 cm and 96 × 96 matrix, resulting in a voxel size of 2.1 × 2.1 × 3mm. Stimuli were presented with Matlab (Psychtoolbox; Brainard, 1997) and displayed on a 40” high definition LCD screen (NordicNeuroLab) viewed by the participants through a mirror located in the scanner. Anatomical MPRAGE images were collected with 1 × 1 × 1 mm resolution, echo time = 2.6ms, TR = 1.75s.

The functional face localizer included two runs. Each run included blocks of dynamic faces and objects. Each block lasted 16 seconds and included 16 stimuli. Category block order was counterbalanced within and across runs. Each localizer run consisted of eight blocks for each category and five blocks of a baseline fixation point resulting in a total of 21 blocks (336s). To maintain vigilance, participants were instructed to press a response box button whenever two identical video-clips appeared consecutively (a 1-back task). This happened twice per block in a random location.

The MT functional localizer included two runs. Each run included blocks of dynamic and static dots. Each block lasted 16s. Category block order was counterbalanced within and across runs. Each localizer run consisted of eight blocks for each category and five blocks of a baseline fixation point resulting in a total of 21 blocks (336s). Participants were instructed to passively view the dots.

The main experiment included four runs. Each run included 21 blocks: 5 baseline fixation blocks and 4 blocks for each of the four experimental conditions: Dynamic/Static x Expression/Gender. Each block lasted 16 seconds and included 3 trials. Each trial consisted of a 4s stimulus (a 4s movie-clip in the dynamic conditions; four images from the movie presented for 1s each in the static conditions) and an inter-trial-interval (ITI) of 1.33 seconds. The order of the experimental conditions was counterbalanced within and across runs. Subjects’ task was to discriminate the gender (male/female) or expression (positive/negative) of each face. During each fixation block, instructions appeared on the screen to inform the subject of the task to be performed on the next block. Subjects responded by pressing one of two response buttons, either while the stimulus was presented or during the ISI. The runs of the localizers and the main experiment were interleaved.

#### Data Analysis

##### Behavioral Data Analysis

We calculated accuracy (percentage of correct responses) and response times for correct responses for each of the four experimental conditions separately. Prior to statistical analysis RT values were converted to log(RT). The behavioral data analysis is based on the results of 12 out of our 18 subjects, as the responses of 6 subjects were not recorded due to technical issues.

##### fMRI Data Analysis

fMRI analysis was performed using statistical parametric mapping (SPM8; http://www.fil.ion.ucl.ac.uk/spm; Wellcome Department of Cognitive Neurology, London, UK)(Acton and Friston, 1998). The first three volumes in each run were acquired during a blank screen display and were discarded from the analysis as “dummy scans”. The data were then preprocessed using slice timing correction and realignment to the first slice of each run. Spatial smoothing was performed for the localizer data only (5mm). A GLM was run with separate regressors for each run and for each condition (i.e. for the face localizer: Faces and Objects; for the MT localizer: Dynamic dots and Static dots; for the main experiment: Dynamic faces-Expression task, Static faces-Expression task, Dynamic faces-Gender task, and Static faces-Gender task).

##### ROI Analysis

For both the face and the MT functional localizer data, category-selective voxels were defined using contrast *t*-maps to assure their functional specificity. Face-selective areas in the fusiform gyrus (FFA), the lateral occipital cortex (OFA), the posterior superior temporal sulcus (pSTS-FA) and the inferior frontal gyrus face area (IFG-FA) were defined based on the dynamic faces > dynamic objects contrast *t*-map (p < .00001, uncorrected). The motion-selective area MT was defined based on the dynamic dots > static dots contrast *t*-map (p < .00001, uncorrected). Because most subjects showed a relatively small MT region using this contrast or no activation at all (n=10), for those subjects who did not show activation we lowered the statistical threshold to p=.01 and used the MarsBaR ‘Draw ROI’ option (Brett, 2002) to define a spherical ROI with a radius of 6mm centered at the peak of the contrast map (average MNI coordinates: right MT (40,-71,12); left MT (-41,-73,9), consistent with previous reports, e.g. Kolster et al., 2010). Table 1 presents the average volume of each ROI (number of voxels) across participants and the number of participants in whom they could be defined.

**Table 1.** The volume (averaged number of voxels and standard deviation) of the functionally defined ROIs and the number of participants^1^ in which each of these areas was found.

|  |  | FFA | OFA | pSTS-FA | MT | IFG-FA |
| --- | --- | --- | --- | --- | --- | --- |
| Experiment 1 | Right | 171 (97)<br>(n=17) | 221<br>(134)<br>(n=10) | 490<br>(368)<br>(n=17) | 134 (94)<br>(n=13) | 90 (87)<br>(n=5) |
|  | Left | 86 (65)<br>(n=16) | 169<br>(231)<br>(n=12) | 324<br>(402)<br>(n=17) | 130<br>(104)<br>(n=13) | 39 (30)<br>(n=4) |
| Experiment 2 | Right | 142<br>(157)<br>(n=16) | 217<br>(145)<br>(n=14) | 487<br>(371)<br>(n=16) | 206<br>(155)<br>(n=15) | 72 (94)<br>(n=6) |
|  | Left | 105<br>(133)<br>(n=13) | 125 (69)<br>(n=14) | 336<br>(280)<br>(n=14) | 193<br>(143)<br>(n=15) | 22 (17)<br>(n=2) |

#### Main Experiment

##### Univariate Analysis

BOLD signal time courses for each of the four experimental conditions were extracted from each ROI using the MarsBaR ROI toolbox of SPM (Brett, 2002). The peak of the BOLD signal was between the third and eighth TRs, which is when the 16s block ended. Accordingly, the dependent measure was an average of the BOLD signal values in TRs 3–8. Statistical analysis of the average response to each of the four conditions within each ROI was performed with Statistica 13.

##### Multivariate Analysis

For the multivariate analysis, we used the β estimates that were created from the GLM used for the univariate analysis (see above). Similarities between voxel-wise patterns of effect-sizes (β estimates) were computed using split-half correlations for each ROI, condition, and run of the main experiment. To allow for the comparison across participants and ROIs, we re-defined ROIs with a fixed-size based on the 60 voxels with the highest localizer contrast *t*-values in each area. ROIs with a smaller number of voxels were excluded from the analysis. This was done to avoid dimensionality effects on classifier performance and prevent potential over fitting (Hastie et al., 2001), while ensuring a sufficient amount of data for effective classification. However, to assure the robustness of the results, similar analyses were also conducted across a range of ROI sizes (30–80 voxels).

Voxel-wise similarity patterns across conditions were computed as follows: first, for each voxel, the mean β value across all conditions was subtracted from each condition-specific β value. This procedure was performed for each run separately. For each condition, the resulting voxel-wise pattern was then averaged across half of the data (two runs) and correlated with the average pattern that was computed based on the other half of the data. These correlations were computed based on every possible split-half of the data and were finally averaged across split-halves to obtain the similarity measure between the two conditions. This procedure was done for each pair of conditions, resulting in a pair-wise correlation matrix reflecting the degree of similarity between the neural patterns evoked by the four experimental conditions.

Next, we used these pair-wise correlations to compute ‘within’ and ‘between’ similarity measures. Specifically, to examine whether a given ROI carries information about Motion, the ‘within’ correlation was computed as the average of the correlation between Dynamic faces-Gender task and Dynamic faces-Expression task, and the correlation between Static faces-Gender task and Static faces-Expression task; and the ‘between’ correlation was computed as the average of the correlation between Dynamic faces-Gender task and Static faces-Expression task, and the correlation between Static faces-Gender task and Dynamic faces-Expression task. To examine whether a given ROI carries information about Task, the ‘within’ correlation was computed as the average of the correlation between Gender task-Dynamic faces and Gender task-Static faces, and the correlation between Expression task-Dynamic faces and Expression task-Static faces; and the ‘between’ correlation was computed as the average of the correlation between Gender task-Dynamic faces and Expression task-Static faces, and the correlation between Gender task-Static faces and Expression task-Dynamic faces.

### Results

#### Behavioral Results

*Accuracy*: We calculated the percentage of correct responses during the gender and expression tasks for the dynamic and the static faces (Table 2). A two-way ANOVA with Motion (Dynamic, Static) and Task (Expression, Gender) as within-subject factors revealed a main effect of Task (F(1,11)=5.16, p=.04, η_p_^2^=.32), indicating better performance for the gender than the expression task. No other effects or interactions were found.

**Table 2.** Mean and standard deviation of accuracy (percent correct) and reaction time in the gender and expression tasks for the dynamic and the static faces in Experiments 1 and 2.

| Experiment 1 |  | Static Faces<br>(in order) | Dynamic Faces |
| --- | --- | --- | --- |
| Accuracy | Gender Task | 87.8 (8.6) | 90.1 (11.0) |
|  | Expression Task | 81.7 (12.0) | 81.5 (12.0) |
| Reaction time | Gender Task | 2.53 (1.34) | 2.54 (1.23) |
|  | Expression Task | 3.32 (0.99) | 3.12 (1.03) |
| Experiment 2 |  | Static Faces<br>(Scrambled order) | Dynamic Faces |
| Accuracy | Gender Task | 90.3 (8.5) | 83.2 (8.1) |
|  | Expression Task | 83.2 (11.2) | 88.7 (6.7) |
| Reaction time | Gender Task | 1.26 (0.26) | 1.75 (0.26) |
|  | Expression Task | 1.75 (0.43) | 1.85 (0.35) |

*Reaction Time*: A two-way ANOVA with Motion (Dynamic, Static) and Task (Expression, Gender) as within-subject factors on reaction time data revealed a main effect of Task (F(1,11)=15.91, p=.002, η_p_^2^=.59) due to faster reaction time on the gender than the expression task and a Task by Motion interaction (F(1,11)=5.28, p=.04, η_p_^2^=.32) (see Table 2).

Overall behavioral results show no motion advantage in the expression or gender tasks. Thus, the current task did not replicate the motion advantage typically found for expression tasks. We will address this issue in Experiment 2.

### Functional MRI

#### Localization of dynamic face areas and motion area MT

The face areas were defined based on a dynamic face localizer that included blocks of dynamic faces and dynamic objects (see Methods). The motion area was defined based on a motion localizer (see Methods). Importantly, the response of area MT to the dynamic faces and objects that were used in our dynamic face localizer was similar indicating no difference in the amount of motion presented in the two categories (see Figure 5).

**Figure 5.**
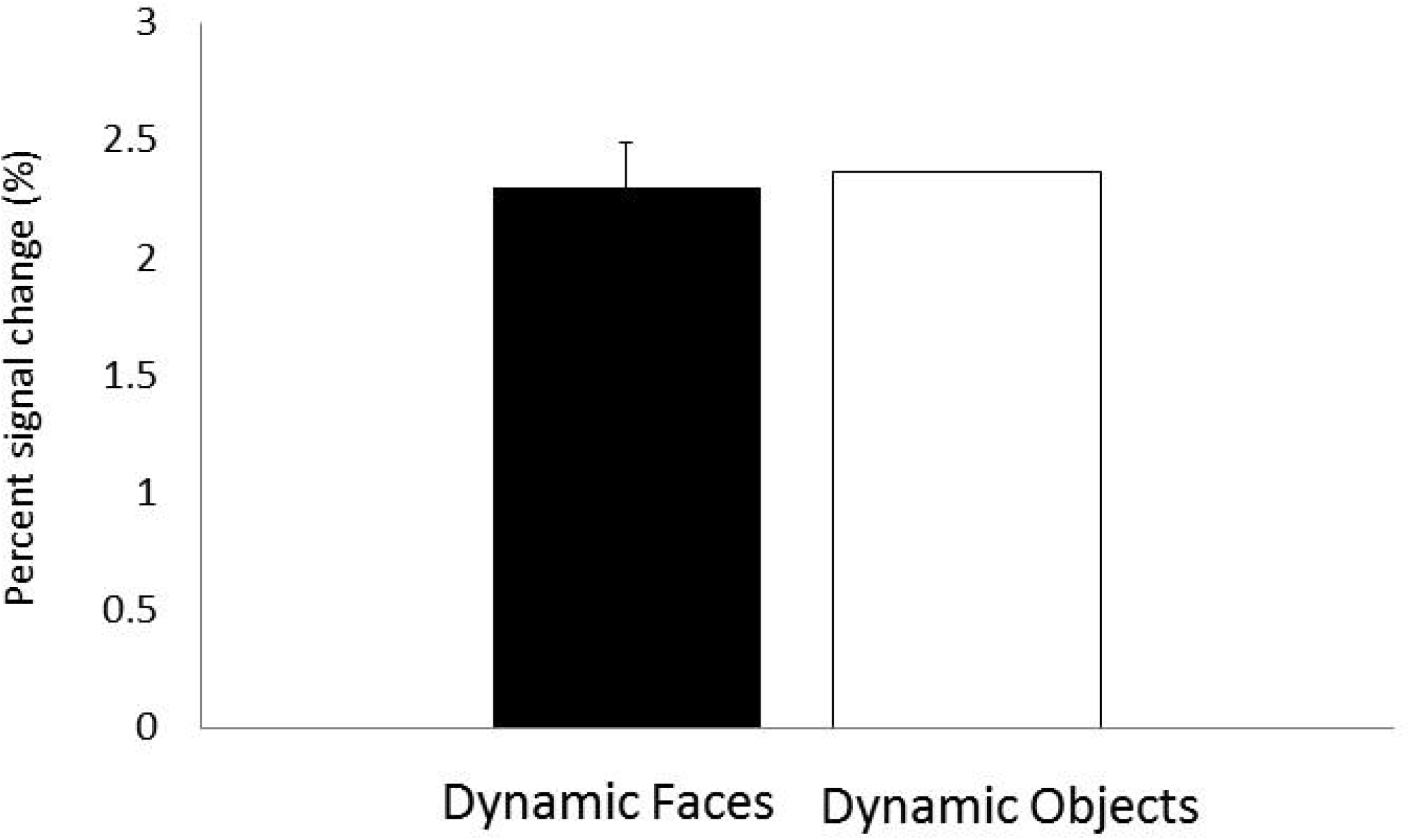
fMRI responses in area MT was similar to the dynamic faces and objects used to localize the face areas. Error bars indicate the standard error of the mean difference between dynamic faces and dynamic objects and are therefore displayed on one of the bars.

Figure 6 shows face-selective and motion areas in a representative subject. Consistent with previous studies, dynamic faces generated a much larger response in the pSTS-FA than in the FFA (see also Table 1). In contrast to most studies that have used static faces and reveal relatively smaller activation in the pSTS-FA (e.g., Kanwisher & Yovel, 2006), the volume of the pSTS-FA was more than twice larger than the FFA and in most subjects extended inferiorly to the lateral occipital sulcus (LOS).

**Figure 6.**
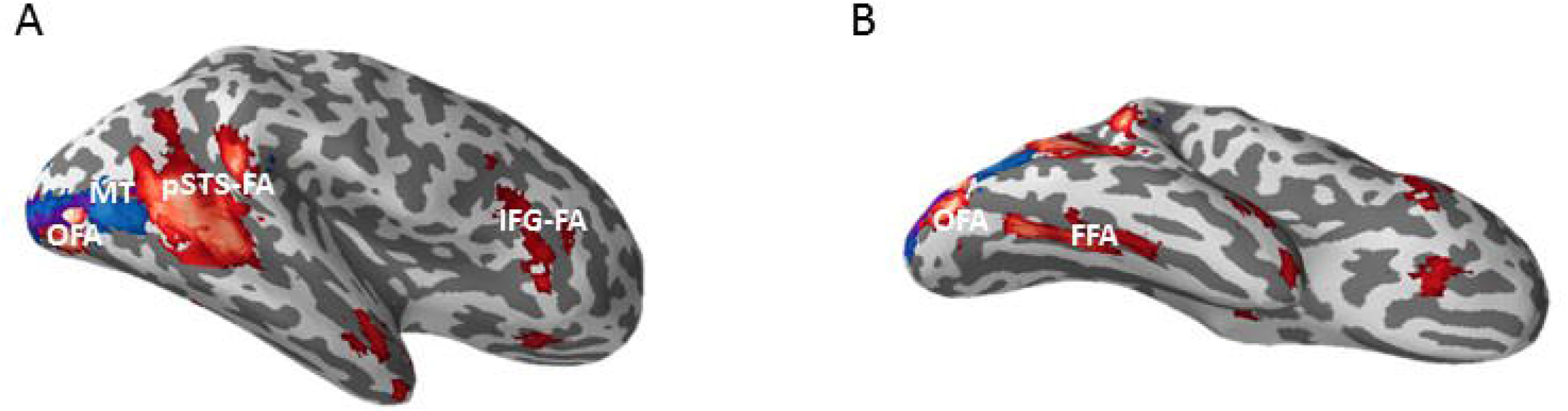
Lateral (A) and ventral (B) views of the right hemisphere of a representative subject showing the face areas, OFA, FFA, pSTS-FA and IFG-FA in red and area MT in blue.

#### The response of the face areas to dynamic and static faces during categorization of changeable and invariant facial aspects

### Univariate analysis

First, we ran an ANOVA with Hemisphere, Motion (Dynamic, Static) and Task (Expression, Gender) as repeated measures to assess whether there was a significant interaction of the effects of interest with hemisphere. This analysis was conducted separately for each ROI (FFA, OFA, pSTS-FA and MT) to maximize the number of subjects that can be included in the analysis. No interactions were found between any of the factors and Hemisphere (p > .1), and therefore for the rest of the analysis, data from the two hemispheres were collapsed using weighted averages based on the volume of each ROI. Because only 5 subjects showed a face-selective response in the right IFG, we report the analysis of data from this region following the results of Experiment 2, where we combined results of the two experiments.

We next examined the effects of the Motion and Task as repeated measures in each ROI (Figure 7, Table 3). The OFA (n= 14) and FFA (n= 17) showed no effects of Motion ((FFA, (F(1,16) < 1, OFA (F(1,13) < 1)) or Task ((FFA, (F(1,16) < 1, OFA (F(1,13) = 2.48, p = .13)) indicating similar engagement during the changeable and invariant tasks and no sensitivity to face motion. In contrast, the pSTS-FA (n= 17) showed a main effect of Motion (F(1,16)=21.19, p=.0002, η_p_^2^=.56) indicating higher response to dynamic than static faces, both during the expression task (t(16)=2.60, p=.01) and the gender task (t(16)=4.71, p=.0002). The pSTS-FA also showed a main effect of task (F(1,16)=4.75, p=.04, η_p_^2^=.22) reflecting higher response during the expression than the gender task. The effect of Task was significant for dynamic faces (t(16)=2.28, p=.03) and marginally significant effect for static faces (t(16)=1.89, p=.07). There was no interaction between Motion and Task (F<1) indicating that the pSTS-FA is sensitive to both facial properties, motion and changeable aspects, independently.

**Figure 7.**
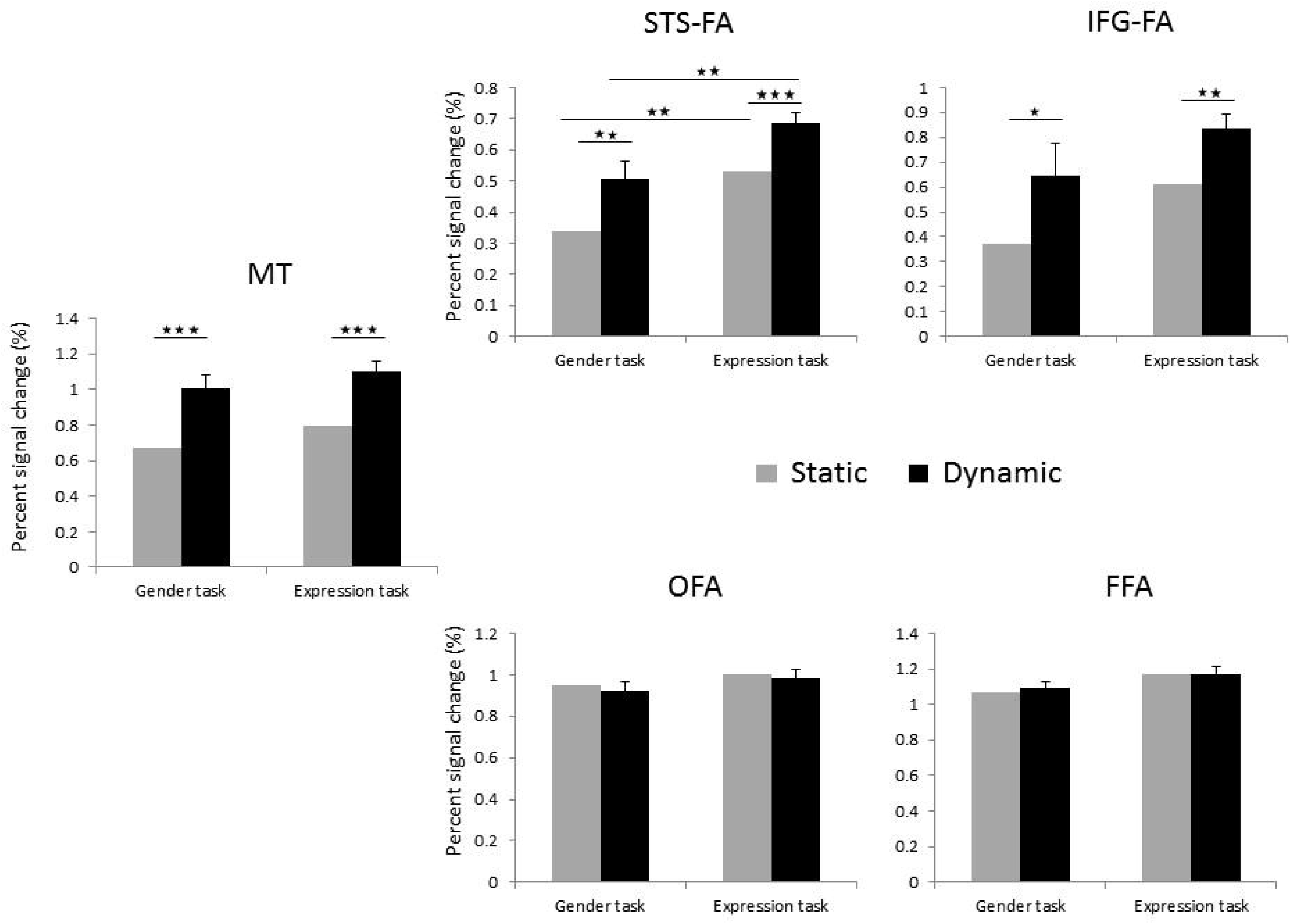
fMRI responses in each ROI including the maximal number of subjects that showed activation in this area (Table 3). Error bars indicate the standard error of the mean difference between static and dynamic faces and are shown on one of the bars. * p<.05; ** p<.01; *** p<.001

**Table 3.** Percent signal change in each ROI for dynamic and static faces during the expression and gender tasks, in Experiments 1 and 2 and the number of subjects (N) that are included in the analysis for each ROI.

|  |  | MT |  | OFA |  | FFA |  | pSTS-FA |  | IFG-FA |  |
| --- | --- | --- | --- | --- | --- | --- | --- | --- | --- | --- | --- |
|  |  | Stat | Dyn | Stat | Dyn | Stat | Dyn | Stat | Dyn | Stat | Dyn |
| Exp. 1 | Gender | 0.62 | 0.93 | 0.90 | 0.83 | 1.00 | 1.02 | 0.36 | 0.51 | 0.43 | 0.41 |
|  | Expression | 0.76 | 1.03 | 1.05 | 1.02 | 1.12 | 1.10 | 0.55 | 0.68 | 0.32 | 0.49 |
|  | N | 14 |  | 14 |  | 17 |  | 17 |  | 5 |  |
| Exp. 2 | Gender | 0.71 | 1.07 | 0.98 | 0.99 | 1.13 | 1.16 | 0.30 | 0.49 | 0.31 | 0.84 |
|  | Expression | 0.82 | 1.16 | 0.96 | 0.94 | 1.20 | 1.22 | 0.50 | 0.68 | 0.85 | 1.12 |
|  | N | 16 |  | 17 |  | 17 |  | 16 |  | 6 |  |

Finally, we examined the response of the motion area MT to the face stimuli during the two tasks. Area MT (n= 14) showed higher response to the dynamic than the static faces (F(1,13)=43.37, p=.00001, η_p_^2^=.76), both during the expression task (t(13)=3.39, p=.004) and during the gender task (t(13)=7.91, p=.0001). No further effects or interactions were found.

To assess whether the patterns of response that we found are significantly different between the dorsal and ventral face areas, we combined the data from the pSTS-FA and FFA and added ROI as a within subject factor in the ANOVA (n=17). This analysis revealed a significant interaction between ROI and Motion (F(1,16)=19.23, p=.0004, η_p_^2^=.54) indicating the sensitivity of the pSTS-FA, but not the FFA, to facial motion. The interaction of ROI and Task (F(1,16) = 2.3, p = .16, η_p_^2^=.13) was not significant indicating that the preference of the pSTS-FA to the expression task was not significantly different from the FFA.

To compare the pattern of response in the OFA and FFA, ANOVA with Motion Task and ROI (OFA, FFA) (n=13) was performed. The analysis did not reveal significant interactions of Motion or Task with ROI (p>.5), indicating that OFA and FFA showed similar pattern of response.

### Multivariate Analysis

Univariate analysis did not reveal effects of motion and task in the ventral face areas, the OFA and FFA. To further assess if information about motion and task is represented in each of the face areas we examined the multi-voxel patterns as a function of motion and categorization task. Split-half correlations between voxel-wise effect sizes were used to assess the degree of similarity between the neural response patterns to the four experimental conditions, within each of the four ROIs. To examine whether a given ROI carries information about Motion, we compared two kinds of correlations: the correlations within presentation type (i.e. correlating data obtained for different dynamic faces, or data obtained for different static faces); to the correlations between presentation types (i.e. correlating data obtained for dynamic faces with data obtained for static faces). If a given brain region is sensitive to motion, the neural patterns generated by two dynamic conditions or by two static conditions should be more similar than the neural patterns generated by a dynamic condition and a static condition. Thus, comparing the correlations within and between dynamic and static stimuli gives a measure of the sensitivity to motion (Haxby et al., 2001, Haxby, 2012).

We first examined these multi-voxel patterns for ROIs of a fixed volume (60 voxels). Higher correlation within presentation type than between presentation types (Dynamic/Static) was found in the pSTS-FA (t(17)=6.70, p=.000004) and in area MT (t(12)=3.67, p=.003) but not in the ventral face areas, the OFA and FFA (p>.3), further supporting the sensitivity to motion in the dorsal but not the ventral face areas, as shown in the univariate results above. The same analysis was conducted using varying sizes of ROIs (30–80 voxels) to verify that the effect is not limited to a specific ROI size. The pSTS-FA and area MT showed higher correlation within than between presentation types across the range of ROI sizes (pSTS-FA: p<.0004 for all ROI sizes; MT: p<.02 for 30, 40, 50 and 60 voxels, p<.08 for 70 and 80 voxels).

To examine whether a given ROI carries information about Task, we compared the correlations within task the expression and gender tasks to the correlations between the two tasks (see Methods). None of the ROIs showed higher correlation within than between tasks (p>.2). Thus, the sensitivity to changeable facial aspects in the pSTS-FA that was found in the univariate analysis is not apparent in the multi-voxel pattern of the neural response of these areas.

### Discussion

Overall, these findings suggest that pSTS-FA is sensitive to motion while the OFA and FFA are not, replicating previous results (Pitcher et al., 2011, 2014). Importantly, we found higher response to motion during both the expression and the gender tasks indicating that the pSTS-FA is primarily sensitive to dynamic information even when subjects performed an invariant facial aspect task. Also consistent with previous findings we revealed that the pSTS-FA showed higher response during the changeable than the invariant facial task for static faces (Hoffman and Haxby, 2000; Engell and Haxby, 2007; Winston et al., 2004). Our findings further show that this effect is also found for dynamic faces. In contrast, the ventral face areas showed no effect of task, consistent with previous findings showing that the FFA is sensitive to facial expression (Ganel et al., 2005; Fox et al., 2009b; Kadosh et al., 2010; Xu & Biederman, 2010; for a recent review see Bernstein and Yovel, 2015). These findings suggest that the pSTS-FA is sensitive to both the dynamic aspects as well as the changeable aspects of faces, possibly due to their dynamic nature, whereas the ventral face areas process the structure of the face regardless of whether it is dynamic or static or the type of facial aspect that is extracted.

In the current study, the static face stimuli were composed of a series of 4 images taken from the movie-clip presented in the order of their appearance. This allowed us to present the richer information that is conveyed by a dynamic face in comparison to a single static face image that was used in previous reports. However, such type of presentation may generate apparent motion for the static condition, which may weaken the effect of motion. Indeed, the behavioral findings that we obtained in this task did not reveal a motion advantage for the expression task as was reported in previous studies that compared performance between dynamic faces and a single static face (for a review see Krumhuber et al., 2013). To examine the difference between the processing of dynamic and static faces while excluding the possible effect of apparent motion, in Experiment 2 the static condition was composed of the same sets of 4 images, presented in a scrambled order.

## Experiment 2

### Materials and Methods

#### Participants

Twenty-one volunteers (11 men, ages 18–32) participated in Experiment 2. Three participants were excluded due to excessive head movements (one participant) or pathological findings (two participants), leaving a total of 18 participants in the final analysis. Participants were paid $15/hr. All participants provided a written informed consent to participate in the study, which was approved by the ethics committees of the Sheba Medical Center and Tel Aviv University.

#### Stimuli

The experiment included three parts: a functional localizer for the face-selective areas, a functional localizer for the motion-selective area MT, and the main experiment. Stimuli of the main experiment were the same as in Experiment 1, with only one difference: in the static condition the same four images that were presented in Experiment 1 in the order of appearance in the movie that they were taken from, were presented in Experiment 2 in a randomly scrambled order. Stimuli of the face functional localizer were exactly as in Experiment 1.

We also changed the stimuli we used to define MT because the moving dots did not generate strong enough activations. In Experiment 2 stimuli of the MT functional localizer were moving concentric grayscale rings, and static shots of these rings (e.g., Weiner & Grill-Spector, 2011).

Procedure and analysis were similar to Experiment 1. The only difference was that subjects were asked to respond once they made a decision, which resulted in shorter reaction times.

### Results

#### Behavioral Results

Accuracy: We calculated the percentage of correct responses during the gender and expression tasks for the dynamic and the static faces. Table 2 summarizes the mean and standard deviation for the different conditions. A two-way ANOVA with Motion (Dynamic, Static) and Task (Expression, Gender) as within-subject factors revealed a main effect of Motion (F(1,17)=6.36, p=.02, η_p_^2^=.27), a main effect of Task (F(1,17)=5.44, p=.03, η_p_^2^=.24), and an interaction between the two factors (F(1,17)=4.51, p=.04, η_p_^2^=.21), indicating that expression recognition was more accurate for dynamic than static faces (t(17)=3.02, p=.007), while in the gender task performance was similar for the dynamic and static faces (p>.3). Thus, the scrambling of the static images resulted in the motion advantage for expression recognition, in line with previous studies that compared expression categorization between dynamic stimuli and one static image (for a review see Krumhuber et al., 2013). As seen below, despite these differences in performance across the two experiments, they both yielded similar pattern of fMRI findings.

Reaction Time: A two-way ANOVA with Motion (Dynamic, Static) and Task (Expression, Gender) as within-subject factors on reaction time revealed a main effect of task due to faster reaction times in the gender than the expression task (F(1,17)=53.02, p=.0001, η_p_^2^=.75) and no effect of Motion nor interaction.

#### fMRI Results

##### Univariate Analysis

We first assessed for each ROI (FFA, OFA, pSTS-FA, and MT) whether there was a significant interaction of the effects of interest with hemisphere. Because we found no interaction of any of the factors of interest with Hemisphere (p > .1), data from the two hemispheres were averaged using weighted averages based on the volume of each ROI. Table 3 shows the percent signal change in each ROI for the four different conditions.

We then assessed the effect of Task and Motion in each ROI using a repeated measures ANOVA. The FFA (n = 17) and OFA (n=17) showed no effect of Task (FFA: F(1,16) < 1, OFA: F(1,16) < 1) or Motion (FFA:F(1,16) < 1, OFA: F(1,16) < 1). In contrast, the pSTS-FA (n = 16) showed a main effect of motion (F(1,15) = 7.36, p < .01, η_p_^2^=.33) due to higher response to dynamic than static faces and a main effect of Task (F(1,15) = 7.22, p < .02, η_p_^2^=.32) indicating higher response during the expression than the gender task. The interaction was not significant. The larger response during the expression task was found for both static (t(15)=2.25, p=.03) and dynamic faces (t(15)=2.03, p=.05). Finally, area MT (n = 16) showed a main effect of Motion (F(1,15) = 13.33, p < .005, η_p_^2^=.47) due to higher response to dynamic than static faces but no effect of Task or an interaction between Task and Motion. These findings replicate results of Experiment 1 indicating that the response of the face and motion areas to dynamic and static faces during a changeable and an invariant task is similar regardless of the order in which the four static face stimuli were presented and the differences in behavioral performance that the two tasks yielded.

Overall, the results of Experiments 1 and 2 revealed similar findings, therefore we combined data from the two experiments to examine the effects on a larger sample. We performed a mixed ANOVA with Experiment as a between subject factor and Task and Motion as within-subject factors. None of the ROIs showed an interaction with Experiment. The Results of the data combined across the two experiments are shown in Figure 7.

The OFA (n = 31) and FFA (n = 34) showed no effects of Task (OFA: F(1,29) = 1.41, p = 25, FFA(1,32) = 2.3, p < .13) or Motion (OFA: F(1,29) < 1, FFA(1,32) < 1). In contrast, the pSTS-FA (n = 33) showed a main effect of Motion (F(1,32)=20.14, p<.0001, η_p_^2^=.39) as well as a main effect of Task (F(1,32)=11.49, p=.002, η_p_^2^=.27) with no interaction between the two factors (F(1,32)=0.04, p=.83). The response to dynamic faces was larger than static faces both during the expression task (t(32)=4.31, p=.00001) and during the gender task (t(32)=3.07, p=.004). The response during the expression task was larger than during the gender task for both dynamic (t(32)=3.08, p=.004) and static faces (t(32)=2.94, p=.006).

Area MT (n= 30) showed higher response to the dynamic than the static faces (F(1,28)=33.58, p=.0001, η_p_^2^=.54), both during the expression task (t(29)=5.65, p=.00001) and during the gender task (t(29)=4.70, p=.00001).

The larger sample allowed us to explore the pattern of response in the inferior frontal gyrus face area (IFG-FA) which was functionally identified in 11 subjects across the two experiments. The IFG-FA showed a main effect of Motion (F(1,9)=57.19, p<.0001, η_p_^2^=.86) indicating a higher response to dynamic than static faces. There was no interaction between Motion and Task indicating that the higher response to dynamic faces was found for both changeable (t(10)=3.84, p=.003) and invariant facial aspects (t(10)=2.16, p=.05). Despite numerically higher response during the expression than the gender task (see Figure 7), the IFG-FA showed no effect of Task (F(1,9) = 1.9, p = .19).

Finally, to examine whether the response of the pSTS-FA was significantly different than the response of the FFA on this larger sample we included ROI (FFA, pSTS-FA) as additional within subject factor. An interaction between ROI and Motion F(1,31) = 31.16, p < .0001, η_p_^2^=.50) indicated greater sensitivity to motion than static faces in the pSTS-FA but not the FFA. An interaction between ROI and Task F(1,31) = 7.68, p < .01, η_p_^2^=.20) indicated greater sensitivity during the expression than the gender task in the pSTS-FA but not the FFA. The 3-way interaction of ROI, Motion and Task was not significant (F(1,31) < 1) indicating that the effect of Task and Motion were additive.

Overall these findings show that the OFA and FFA showed no sensitivity to motion and respond similarly during the expression the gender tasks. The pSTS-FA showed higher response to dynamic than static faces during both the expression and the gender tasks and higher response during the expression than the gender task for both dynamic and static faces. The IFG was also highly responsive to dynamic stimuli but did not show a significantly higher response during the expression than the gender task.

##### Multivariate Analysis

As in experiment 1, split-half correlations were used to assess the degree of similarity between the neural patterns generated by the four experimental conditions, and correlations within vs. between presentation type or task were examined as a measure of sensitivity to motion or task, respectively. Higher correlation within presentation type than between presentation types was found in the pSTS-FA (t(15)=3.41, p=.003) and in area MT (t(12)=3.01, p=.01) but not in the OFA and FFA (p>.2). The pSTS-FA and area MT showed higher correlation within than between presentation types across a range of ROI sizes (30-80 voxels) (pSTS-FA: p<.01 for all ROI sizes; MT: p<.02 for all ROI sizes). None of the ROIs showed higher correlation within than between tasks (p>.15), indicating that they did not carry information about the task that was being performed.

A combined analysis across the two experiments of the multivariate analysis was first examined for ROIs of a fixed volume (60 voxels). Results of data combined from the two experiments are shown in Figure 8. Higher correlation within presentation type (Dynamic/Static) than between presentation types was found in the pSTS-FA (t(33)=4.3, p=.0001), IFG-FA (t(7)=3.61, p=.008) and in area MT (t(22)=4.77, p=.000) but not in the OFA and FFA (p>.3), further supporting the sensitivity to motion in the dorsal but not the ventral face areas, as shown in the univariate results reported above. The same analysis was conducted using varying ROI volumes (30–80 voxels) to verify that the effect is not limited to a specific ROI size. The pSTS-FA, IFG-FA and area MT showed higher correlation within than between presentation types across the range of ROI sizes (pSTS-FA: p<.0004 for all ROI sizes; IFG-FA: p<.04 for all ROI sizes; MT: p<.02 for 30, 40, 50 and 60 voxels, p<.08 for 70 and 80 voxels). We then compared the correlations within and between Tasks. None of the ROIs showed higher correlation within than between tasks (p>.2) (see Figure 8). Thus, the sensitivity to changeable facial aspects in the pSTS-FA in the univariate analysis is not apparent in the multi-voxel pattern of the neural response of these areas.

**Figure 8.**
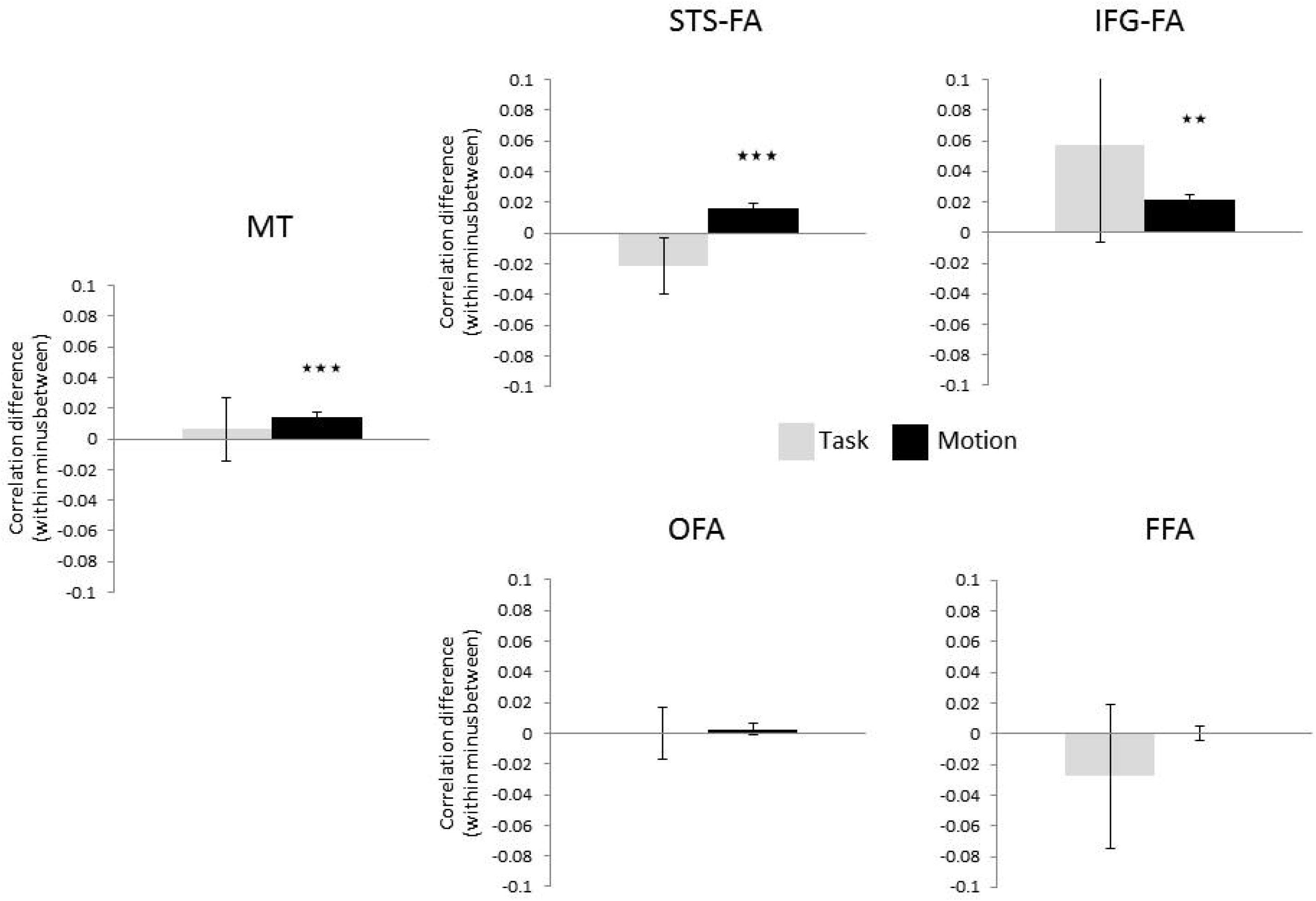
Pattern similarity analysis suggests sensitivity to motion in the pSTS-FA, IFG-FA and area MT but not in the OFA and FFA. Error bars indicate the standard error of the correlation difference ** p < .01; *** p<.001.

To assess whether the sensitivity to motion based on pattern analysis in the dorsal and ventral face areas is significantly different, we performed t-tests to directly compare data from the pSTS-FA and FFA. We found that the sensitivity to motion was significantly larger in pSTS-FA than in the FFA (t(20)=3.07, p=.006). The two ventral face areas, OFA and FFA, were similarly insensitive to motion (p>.6), and the two dorsal face areas, pSTS-FA and IFG-FA, were similarly sensitive to motion (p>.4).

### Functional Connectivity

Results so far suggest functional dissociation between the pSTS-FA and the ventral face areas, OFA and FFA. These findings may be consistent with previous anatomical connectivity (Gschwind et al., 2012, Pyles et al., 2013) and functional correlation (Avidan et al., 2014) studies that reported stronger structural connections and synchronization between OFA and FFA than with the pSTS-FA. Here, we were interested in replicating these findings as well as examining the functional connectivity between the face areas and the motion area MT. This allowed us to evaluate the model suggested by O’Toole and colleagues according to which area MT is connected with both the pSTS-FA for the processing of dynamic information as well as with the ventral face areas for the extraction of form from motion (O’Toole et al., 2002) (Figure 1B). Therefore, we examined the functional synchronization between the face areas and MT by correlating their respective signal time-courses recorded during the main task.

To assess the relative strength of functional connections between the different ROIs, the correlations were converted to Fisher Z scores and a repeated-measures ANOVA with ROI pairs (MT-OFA, MT-FFA, MT-pSTS-FA, OFA-FFA, OFA-pSTS-FA, FFA-pSTS-FA) and Hemisphere was performed. The ANOVA revealed a significant effect of ROI pairs (F(5,60)=12.60, p=.0000, η_p_^2^=.51) and a significant interaction of ROI pairs and Hemisphere (F(5,60)=3.18, p=.01, η_p_^2^=.20), therefore correlations are reported separately for the right and left ROIs. As shown in Figure 9A, the OFA was strongly correlated with the FFA but less correlated with the pSTS-FA (right: t(91)=12.19, p=.000; left: t(91)=14.95, p=.000). Area MT was similarly correlated with all other ROIs (p>.3).

**Figure 9.**
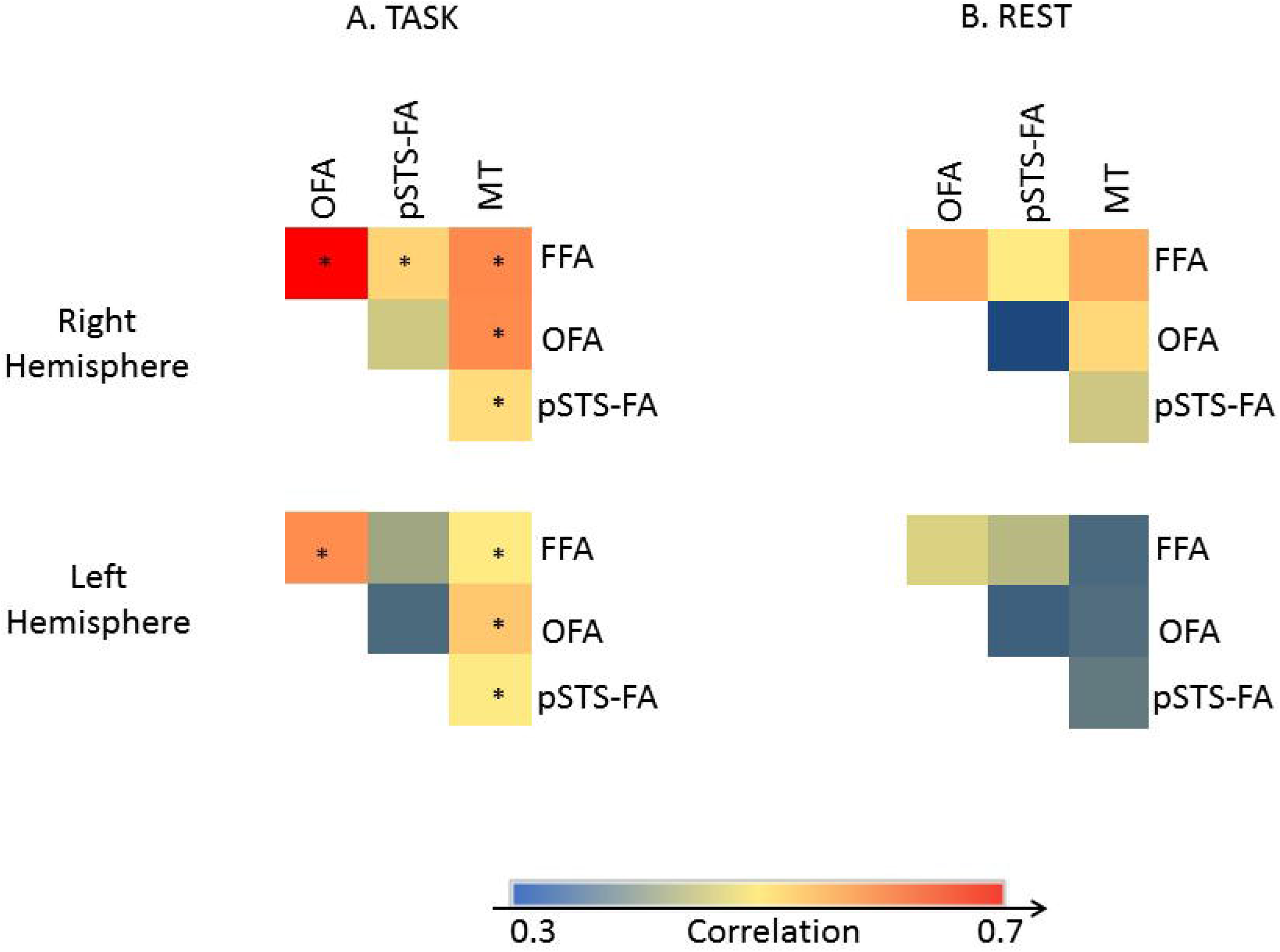
Functional connectivity of the face processing system during task (A) and rest (B). The color code indicates the level of correlation calculated between each pair of regions in each subject and then averaged across subjects. * p < .05

To ensure that these correlation patterns are independent of task, we examined the functional synchronization across the face-processing network during “rest”. In addition to the dynamic face localizer and motion localizer, additional 12 subjects underwent a 10-minute resting-state fMRI scan in which they were instructed to close their eyes and let their mind wander without falling asleep. Functional correlations were then computed between the BOLD-signal time-courses of the different ROIs, which were functionally localized using the same localizers described above.

Figure 9B presents matrices of pairwise correlations between the face-selective areas and area MT, averaged across subjects. An ANOVA with ROI pairs and Hemisphere as repeated measures and Type of processing (Face processing task, Rest) as a between-subjects factor revealed a significant effect of ROI pairs (F(5,85)=9.31, p=.0000, η_p_^2^=.35) but no interactions with the Type of processing (p>.4), indicating that the functional correlation patterns were similar across rest and task. Thus, the resting state functional correlations showed similar patterns of strong synchronization between the ventral areas FFA and OFA, weaker synchronization between the OFA and the pSTS-FA, and synchronization of all three regions with area MT.

### General Discussion

The goal of the current study was the integrate two major functional characteristics of the ventral and dorsal face areas, their sensitivity to changeable vs. invariant facial aspects (Haxby et al., 2000) and their different sensitivity to motion (O’Toole et al., 2002, Pitcher et al., 2011, 2014). Our results show that the dorsal face areas, pSTS-FA and IFG-FA responded more strongly to dynamic than to static faces (Fox et al., 2009a, Pitcher et al., 2011) regardless of whether participants performed an expression or a gender task. The pSTS-FA also responded more strongly during the expression than the gender task regardless of whether faces were moving or not. In contrast, the ventral face areas, FFA and OFA, were responsive to faces regardless of whether they were moving or not (consistent with (Fox et al., 2009a, Pitcher et al., 2011) and whether subjects categorize invariant or changeable facial aspects (for a recent review see (Bernstein and Yovel, 2015) (Figure 7). These findings critically constrain the relationship between the processing of changeable facial aspects and the processing of facial motion. First, the processing of motion cannot be reduced to the processing of changeable aspects: even when the task requires categorizing invariant aspects, the dorsal face areas are still engaged more strongly in processing dynamic vs. static faces. Thus, this finding is inconsistent with a model that dissociates the processing of changeable and invariant face aspects in the dorsal and ventral face pathways, respectively (Haxby et al., 2000, Gobbini and Haxby, 2007) (Figure 3A). Second, the processing of changeable aspects cannot be fully accounted for by the processing of facial motion: even in the absence of such motion, the pSTS-FA is still engaged more strongly by processing changeable vs. invariant aspects. More generally, whereas the primary division between the dorsal and ventral face areas appears to separate face motion from form (O’Toole et al., 2002, Pitcher et al., 2014) (Figure 3B), this dissociation alone cannot exclusively account for the functional differences between the two face pathways.

Still, processing dynamic information and processing changeable aspects are not entirely dissociated and appear to be integrated within the face-processing network: regions within this network are either sensitive to both (pSTS-FA) or to neither (OFA and FFA). Specifically, we suggest that the preferential response to changeable facial aspects in the pSTS-FA may be due to their dynamic nature, given that the processing of changeable aspects such as expression relies on dynamic changes of facial structure over time. The association between changeable facial aspects and motion is supported by a large body of behavioral evidence showing a dynamic advantage for expression recognition (for a review see (Krumhuber et al., 2013), contrasting with a less conclusive dynamic advantage for identity recognition (for a review see (O’Toole, 2010).

The integrated neural framework of face processing proposed here (Bernstein and Yovel, 2015, Duchaine and Yovel, 2015) (Figure 2) is consistent with several reports that have been recently published and may not easily fit with current models. First, whereas the classic models of face recognition (Bruce and Young, 1986, Haxby et al., 2000) proposed parallel and independent routes for invariant and changeable facial aspects, recent studies have found that the processing of identity and expression may be mediated by the same system rather than being fully separated (for a review see (Lander and Butcher, 2015). A common mechanism for the processing of changeable and invariant aspects is also supported by computational and neuropsychological data (Calder and Young, 2005), and by fMR-adaptation studies showing sensitivity of the FFA to both facial expression and identity (Ganel et al., 2005, Fox et al., 2009b, Kadosh et al., 2010, Xu and Biederman, 2010). These findings suggest that the primary division of labor between the ventral and dorsal face areas may not be the division to invariant and changeable aspects.

Second, whereas the current dominant model (Haxby et al., 2000, Gobbini and Haxby, 2007) posits that the OFA is a source of input to both the ventral and the dorsal face pathways, recent connectivity studies (Gschwind et al., 2012, Pyles et al., 2013) have instead revealed that the OFA may be connected to the FFA more than it is to the pSTS-FA. In line with previous work (Avidan et al., 2014), we found stronger functional correlation between the OFA and FFA and weaker correlation with the pSTS-FA, both during face processing tasks and at rest. These findings are in line with the similar response profiles to face stimuli in the FFA and OFA, which differ from the response profile of the pSTS-FA. O’Toole et al (O’Toole et al., 2002) have suggested that the pSTS-FA may receive input from area MT, which is consistent with the finding that both these regions are engaged in motion processing. Indeed, we found functional correlations between area MT and the pSTS-FA, although area MT was also correlated with the FFA and OFA. We suggest that these latter correlations between area MT and the ventral face areas may allow for “form-from-motion” analysis, as proposed by O’Toole and colleagues (O’Toole et al., 2002). According to their model, structural information is extracted from dynamic faces in area MT and sent back to the ventral stream as ‘motionless’ form information for structural analysis. Importantly, this is the first study to our knowledge that specifically examined functional coupling between area MT and the face selective areas, thereby providing empirical support for the connections between them as proposed by O’Toole and colleagues. It is noteworthy that the face areas in the current study were defined using a dynamic face localizer, which may result in higher functional correlations between all face areas and area MT relative to areas defined by static images. However, given the large overlap between the ventral areas that are identified using static and dynamic functional localizers (Fox et al., 2009a), the patterns of their functional synchronization with MT are likely to be similar. Alternatively, the current functional correlations specifically reflect association of the motion area with the processing of dynamic faces by the face network.

In addition to the face-selective discussed above, recent studies that used a dynamic face localizer have reported a prefrontal face-selective area, typically located in the IFG (Fox et al., 2009a, Pitcher et al., 2011). The functional profile of the IFG face area (IFG-FA) has been only scarcely studied (for a review see (Chan, 2013), possibly due to its relatively low response to the prevalently used static face stimuli. Here, we functionally defined this region with a dynamic face localizer and still found it only in less than 1/3 of the subjects we scanned (see also Pitcher et al., 2011). Similar to the pSTS-FA, the IFG-FA showed a stronger response to dynamic than static faces. Unlike the pSTS-FA, the response of the IFG-FA was not statistically significant greater during the expression than the gender task, despite a large numerical difference (see Figure 7), which may be due to the low power of the analysis. Nevertheless, recent studies did indicate that the IFG-FA may play a role in the processing of eye-gaze information (Chan and Downing, 2011). However, the IFG-FA was shown to be involved also in the representation of face identity (Guntupalli, 2016). Thus, the precise functional role of the IFG-FA remains to be elucidated in future studies. Moreover, the IFG-FA may play a role not only in perception but also in action: it has been identified in studies that examined the human mirror-neuron system as being responsive to both viewed and performed motion (Kilner et al., 2009, Molenberghs et al., 2012). Future studies will assess the extent to which the response of the IFG-FA to dynamic faces is purely perceptual or whether the face-selective activation in the IFG may be part of the mirror system and therefore respond also to face and body motor movements and to dynamic body stimuli.

Finally, the dorsal face areas may be part of a more general system for the processing of the whole dynamic person (for review see (Yovel and O’Toole, 2016). In particular, the STS has been shown to be activated by dynamic bodies such as point light displays (Allison et al., 2000, Beauchamp et al., 2003, Grossman et al., 2005), dynamic full light bodies (Hahn and O’Toole, 2016) as well as voices (Belin et al., 2000, Watson et al., 2014). Thus, the STS is considered a neural hub for multi-modal representation of the whole dynamic person.

In summary, we report a comprehensive and systematic characterization of two central functional characteristics of the face processing network that have so far studied separately - sensitivity to changeable and invariant facial aspects and to dynamic and static facial stimuli. Our revised framework goes beyond previous accounts in that it shows that (i) the dorsal face area, the pSTS-FA is sensitive to motion and to changeable facial aspects (ii) the ventral face areas, the OFA and FFA, are similarly sensitive to changeable and invariant facial aspects and are insensitive to motion, and (iii) the motion area MT is functionally associated with both the ventral and dorsal dynamic face areas. This integrated model provides an ecologically motivated framework for the functional architecture of the face-processing network, one that can better accommodate the dynamic faces we encounter in the real world.

## Acknowledgements

This work was supported by a grant from the Israeli Science Foundation ISF 392/16 to GY, and a travel fellowship to MB from the Daniel Turnberg Travel Fellowship Scheme for shortterm exchange visits between the UK and the Middle East.

## Footnotes

1 Note that the number of subjects in each ROI may not reflect the number of subjects that included in the analysis in which data from the two hemispheres were averaged (see Data Analysis). For example, for the OFA in Experiment 1, the table indicates 10 and 12 subjects in right and left OFA, respectively, but overall 14 subjects were included in the analysis because 8 subjects showed activations in both hemispheres, 2 subjects showed only right hemisphere activation and 4 subjects showed only left hemisphere activations resulting in 14 subjects across both hemispheres.

## References

Acton PD, Friston KJ 1998 Statistical parametric mapping in functional neuroimaging: beyond PET and fMRI activation studies. European journal of nuclear medicine 25:663–667.

Allison T, Puce A, McCarthy G 2000 Social perception from visual cues: role of the STS region. Trends Cogn Sci 4:267–278.

Ambadar Z, Schooler JW, Cohn JF 2005 Deciphering the enigmatic face - The importance of facial dynamics in interpreting subtle facial expressions. Psychol Sci 16:403–410.

Avidan G, Tanzer M, Hadj-Bouziane F, Liu N, Ungerleider LG, Behrmann M 2014 Selective Dissociation Between Core and Extended Regions of the Face Processing Network in Congenital Prosopagnosia. Cerebral cortex 24:1565–1578.

Axelrod V, Yovel G 2015 Successful Decoding of Famous Faces in the Fusiform Face Area. Plos One 10.

Baseler HA, Harris RJ, Young AW, Andrews TJ 2013 Neural responses to expression and gaze in the posterior superior temporal sulcus interact with facial identity. Cerebral cortex 24:737–744.

Bassili JN 1978 Facial Motion in Perception of Faces and of Emotional Expression. J Exp Psychol Human 4:373–379.

Bassili JN 1979 Emotion Recognition - Role of Facial Movement and the Relative Importance of Upper and Lower Areas of the Face. J Pers Soc Psychol 37:2049–2058.

Beauchamp MS, Lee KE, Haxby JV, Martin A 2003 fMRI responses to video and point-light displays of moving humans and manipulable objects. J Cognitive Neurosci 15:991–1001.

Belin P, Zatorre RJ, Lafaille P, Ahad P, Pike B 2000 Voice-selective areas in human auditory cortex. Nature 403:309–312.

Bernstein M, Yovel G 2015 Two neural pathways of face processing: A critical evaluation of current models. Neurosci Biobehav R 55:536–546.

Blank I, Kanwisher N, Fedorenko E 2014 A functional dissociation between language and multiple-demand systems revealed in patterns of BOLD signal fluctuations. Journal of neurophysiology 112:1105–1118.

Brett MA, J.; Valabregue, R.; Poline, J. 2002 Region of interest analysis using an SPM toolbox. Neuroimage 16.

Bruce V, Young A 1986 Understanding face recognition. British journal of psychology 77 Pt 3:305–327.

Calder AJ, Young AW 2005 Understanding the recognition of facial identity and facial expression. Nature reviews Neuroscience 6:641–651.

Ceccarini F, Caudek C 2013 Anger superiority effect: The importance of dynamic emotional facial expressions. Vis Cogn 21:498–540.

Chan AW 2013 Functional organization and visual representations of human ventral lateral prefrontal cortex. Front Psychol 4:371.

Chan AW, Downing PE 2011 Faces and eyes in human lateral prefrontal cortex. Frontiers in human neuroscience 5:51.

Cordes D, Haughton VM, Arfanakis K, Carew JD, Turski PA, Moritz CH, Quigley MA, Meyerand ME 2001 Frequencies contributing to functional connectivity in the cerebral cortex in “resting-state” data. AJNR American journal of neuroradiology 22:1326–1333.

Duchaine B, Yovel G 2015 A Revised Neural Framework for Face Processing. Annu Rev Vis Sc 1:393–416.

Engell AD, Haxby JV 2007 Facial expression and gaze-direction in human superior temporal sulcus. Neuropsychologia 45:3234–3241.

Fisher C, Freiwald WA 2015 Contrasting Specializations for Facial Motion within the Macaque Face-Processing System. Curr Biol 25:261–266.

Fox CJ, Iaria G, Barton JJS 2009a Defining the Face Processing Network: Optimization of the Functional Localizer in fMRI. Hum Brain Mapp 30:1637–1651.

Fox CJ, Moon SY, Iaria G, Barton JJ 2009b The correlates of subjective perception of identity and expression in the face network: an fMRI adaptation study. Neuroimage 44:569–580.

Freiwald W, Duchaine B, Yovel G 2016 Face Processing Systems: From Neurons to Real-World Social Perception. Annu Rev Neurosci 39:325–346.

Fujimura T, Suzuki N 2010 Effects of dynamic information in recognising facial expressions on dimensional and categorical judgments. Perception 39:543–552.

Furl N, van Rijsbergen NJ, Treves A, Friston KJ, Dolan RJ 2007 Experience-dependent coding of facial expression in superior temporal sulcus. Proceedings of the National Academy of Sciences of the United States of America 104:13485–13489.

Ganel T, Valyear KF, Goshen-Gottstein Y, Goodale MA 2005 The involvement of the “fusiform face area” in processing facial expression. Neuropsychologia 43:1645–1654.

Gilaie-Dotan S, Malach R 2007 Sub-exemplar shape tuning in human face-related areas. Cerebral cortex 17:325–338.

Gobbini Ml, Haxby JV 2007 Neural systems for recognition of familiar faces. Neuropsychologia 45:32–41.

Goesaert E, Op de Beeck HP 2013 Representations of facial identity information in the ventral visual stream investigated with multivoxel pattern analyses. The Journal of neuroscience : the official journal of the Society for Neuroscience 33:8549–8558.

Grill-Spector K, Knouf N, Kanwisher N 2004 The fusiform face area subserves face perception, not generic within-category identification. Nat Neurosci 7:555–562.

Grossman ED, Battelli L, Pascual-Leone A 2005 Repetitive TMS over posterior STS disrupts perception of biological motion. Vision Res 45:2847–2853.

Gschwind M, Pourtois G, Schwartz S, de Ville DV, Vuilleumier P 2012 White-Matter Connectivity between Face-Responsive Regions in the Human Brain. Cerebral cortex 22:1564–1576.

Guntupalli JSW, K. G.; Gobbini, I. 2016 Disentangling the representation of identity from head view along the human face processing pathway.

Hahn CA, O’Toole AJ 2016 Recognizing Approaching Walkers: Neural Decoding of Person Familiarity in Cortical Areas Responsive to Faces, Bodies, and Biological Motion. Neuroimage.

Hastie T, Tibshirani R, Friedman JH 2001 The elements of statistical learning : data mining, inference, and prediction : with 200 full-color illustrations. New York: Springer.

Haxby JV 2012 Multivariate pattern analysis of fMRI: The early beginnings. Neuroimage 62:852–855.

Haxby JV, Gobbini MI, Furey ML, Ishai A, Schouten JL, Pietrini P 2001 Distributed and overlapping representations of faces and objects in ventral temporal cortex. Science 293:2425–2430.

Haxby JV, Hoffman EA, Gobbini MI 2000 The distributed human neural system for face perception. Trends Cogn Sci 4:223–233.

Huk AC, Dougherty RF, Heeger DJ 2002 Retinotopy and functional subdivision of human areas MT and MST. The Journal of neuroscience : the official journal of the Society for Neuroscience 22:7195–7205.

Kadosh KC, Henson RNA, Kadosh RC, Johnson MH, Dick F 2010 Task-dependent Activation of Face-sensitive Cortex: An fMRI Adaptation Study. J Cognitive Neurosci 22:903–917.

Kanwisher N, Yovel G 2006 The fusiform face area: a cortical region specialized for the perception of faces. Philos T R Soc B 361:2109–2128.

Kaulard K, Cunningham DW, Bulthoff HH, Wallraven C 2012 The MPI Facial Expression Database - A Validated Database of Emotional and Conversational Facial Expressions. Plos One 7.

Kilner JM, Neal A, Weiskopf N, Friston KJ, Frith CD 2009 Evidence of mirror neurons in human inferior frontal gyrus. The Journal of neuroscience : the official journal of the Society for Neuroscience 29:10153–10159.

Kolster H, Peeters R, Orban GA 2010 The retinotopic organization of the human middle temporal area MT/V5 and its cortical neighbors. The Journal of neuroscience : the official journal of the Society for Neuroscience 30:9801–9820.

Krumhuber EG, Kappas A, Manstead ASR 2013 Effects of Dynamic Aspects of Facial Expressions: A Review. Emot Rev 5:41–46.

Lander K, Butcher N 2015 Independence of face identity and expression processing: exploring the role of motion. Front Psychol 6.

Lander K, Davies R 2007 Exploring the role of characteristic motion when learning new faces. Quarterly journal of experimental psychology 60:519–526.

Molenberghs P, Cunnington R, Mattingley JB 2012 Brain regions with mirror properties: a meta-analysis of 125 human fMRI studies. Neurosci Biobehav Rev 36:341–349.

Nestor A, Plaut DC, Behrmann M 2011 Unraveling the distributed neural code of facial identity through spatiotemporal pattern analysis. Proceedings of the National Academy of Sciences of the United States of America 108:9998–10003.

Nusseck M, Cunningham DW, Wallraven C, Bulthoff HH 2008 The contribution of different facial regions to the recognition of conversational expressions. J Vision 8.

O’Toole AJ, Harms J, Snow SL, Hurst DR, Pappas MR, Ayyad JH, Abdi H 2005 A video database of moving faces and people, leee T Pattern Anal 27:812–816.

O’Toole AJ, Roark DA, Abdi H 2002 Recognizing moving faces: a psychological and neural synthesis. Trends Cogn Sci 6:261–266.

O’Toole AJR, D. 2010 Memory for Moving Faces: The Interplay of Two Recognition Systems. In: Dynamic Faces: Insights from Experiments and Computation Curio, C. B., H. H.; Giese, M. A., ed: MIT Press.

O’Toole, A. J., Phillips, P. J., Weimer, S., Roark, D. A., Ayyad, J., Barwick, R., & Dunlop, J. 2011. Recognizing people from dynamic and static faces and bodies: Dissecting identity with a fusion approach. Vision Research, 511, 74–83.

Pitcher D, Dilks DD, Saxe RR, Triantafyllou C, Kanwisher N 2011 Differential selectivity for dynamic versus static information in face-selective cortical regions. Neuroimage 56:2356–2363.

Pitcher D, Duchaine B, Walsh V 2014 Combined TMS and FMRI reveal dissociable cortical pathways for dynamic and static face perception. Curr Biol 24:2066–2070.

Pyles JA, Verstynen TD, Schneider W, Tarr MJ 2013 Explicating the Face Perception Network with White Matter Connectivity. Plos One 8.

Roark DA, O’Toole AJ, Abdi H, Barrett SE 2006 Learning the moves: the effect of familiarity and facial motion on person recognition across large changes in viewing format. Perception 35:761–773.

Rotshtein P, Henson RNA, Treves A, Driver J, Dolan RJ 2005 Morphing Marilyn into Maggie dissociates physical and identity face representations in the brain. Nat Neurosci 8:107–113.

Said CP, Moore CD, Engell AD, Todorov A, Haxby JV 2010 Distributed representations of dynamic facial expressions in the superior temporal sulcus. J Vision 10.

Schultz J, Pilz KS 2009 Natural facial motion enhances cortical responses to faces. Exp Brain Res 194:465–475.

Trautmann SA, Fehr T, Herrmann M 2009 Emotions in motion: Dynamic compared to static facial expressions of disgust and happiness reveal more widespread emotion-specific activations. Brain Res 1284:100–115.

Watson R, Latinus M, Noguchi T, Garrod O, Crabbe F, Belin P 2014 Crossmodal adaptation in right posterior superior temporal sulcus during face-voice emotional integration. The Journal of neuroscience : the official journal of the Society for Neuroscience 34:6813–6821.

Whitfield-Gabrieli S, Nieto-Castanon A 2012 Conn: a functional connectivity toolbox for correlated and anticorrelated brain networks. Brain connectivity 2:125–141.

Xiao NQG, Perrotta S, Quinn PC, Wang Z, Sun YHP, Lee K 2014 On the facilitative effects of face motion on face recognition and its development. Front Psychol 5.

Xu X, Biederman I 2010 Loci of the release from fMRI adaptation for changes in facial expression, identity, and viewpoint. J Vis 10.

Yovel G, Freiwald WA 2013 Face recognition systems in monkey and human: are they the same thing? F1000prime reports 5:10.

Yovel G, Kanwisher N 2005 The neural basis of the behavioral face-inversion effect. Curr Biol 15:2256–2262.

Yovel G, O’Toole AJ 2016 Recognizing People in Motion. Trends Cogn Sci 20:383–395.

